# Gap Junctions Amplify Spatial Variations in Cell Volume in Proliferating Solid Tumors

**DOI:** 10.1101/2020.04.15.043802

**Authors:** Eoin McEvoy, Yulong Han, Ming Guo, Vivek B. Shenoy

## Abstract

Cancer progression is driven by cell proliferation, apoptosis, and matrix invasion, which in turn depend on a myriad of factors including microenvironment stiffness, nutrient supply, and intercellular communication. Cell proliferation is regulated by volume, but in 3D clusters it remains unclear how multiple cells interact to control their size. In this study, we propose a mechano-osmotic model to investigate the evolution of volume dynamics within multicellular systems. Volume control depends on an interplay between multiple cellular constituents, including gap junctions, mechanosensitive ion channels, energy consuming ion transporters, and the actomyosin cortex, that coordinate to manipulate cellular osmolarity. In connected cells, mechanical loading is shown to significantly affect how these components cooperate to transport ions, and precise volume control is impacted by the emergence of osmotic pressure gradients between cells. Consequent increases in cellular ion concentrations drive swelling, while a loss of ions impedes the compression resistance of cells. Combining the modeling framework with novel experiments, we identify how gap junctions can amplify spatial variations in cell volume within multicellular spheroids and, further, describe how the process depends on proliferation-induced solid stress. Our model provides new insight into the role of gap junctions in cancer progression and can help guide the development of therapeutics that target inter- and extra-cellular ion transport.

## Introduction

Cell volume is typically tightly regulated to sustain normal function and survival^1^. In single cells, volume control involves an interplay between ion channels on the membrane that permit passive exchange between the cytosol and extracellular fluid and active ion transporters that move solutes against a concentration gradient^2,3^. As water movement across the cell membrane is largely driven by osmotic pressure, precise control of the cytosolic ion concentration can increase or decrease cell volume. Impairment of ion regulation has severe consequences and is indicated in many disease states; for example, sickle cell dehydration is associated with a pathological loss of erythrocyte ions^4^, and potassium channels play a role in cells resisting apoptosis during cancer development^5^. In multicellular systems, cells adhere to one another via cadherin and catenin mediated complexes^6^. Alongside these adhesions, connexin structures assemble to form gap junctions that permit exchange of ions and fluid between cells^7^. Although these junctions are particularly known to be of importance during development^8^ and in cardiac conductance^9^, they are present in the majority of mammalian cell types. However, despite clear intuition that water and ion flow across gap junctions should confound cytosolic osmolarity and influence cellular shrinkage and swelling, their role in volume dynamics has not been well studied.

In exploring cell behavior, multicellular spheroids have emerged as an increasingly promising experimental model that aim to bridge the gap between in-vitro and in-vivo conditions^10^. Recently, we seeded mammary epithelial cells in hydrogel to investigate how individual cell volumes vary spatially in a proliferating cluster^11^. We identified that peripheral cells became more swollen as the cluster grew, and cells at the core were highly compressed. Blocking gap junctions normalized volume distributions, indicating that they play an important role in mediating differential cell swelling. The development and progression of cancer is driven by a myriad of factors, including matrix density^12,13^, cell adhesion^14,15^, and interactions between different cell types^16,17^, that collectively influence cell proliferation, apoptosis, and matrix invasion. Furthermore, proliferation also depends on cellular size^18,19^ and stress^20,21^. However, in 3D clusters it remains unclear how individual cells coordinate to regulate their volumes.

Motivated by early work on water movement across lipid membranes^22^, a number of analytical models have sought to address how cells regulate their volume via ion exchange with their microenvironment^23–25^. Jiang and Sun (2013)^26^ considered the critical role of cell mechanics and ion channel mechano-sensitivity, highlighting how cells can maintain their volume in response to osmotic shock. Beyond, similar models have been proposed to understand how ion transport governs the swelling or shrinkage of adhered single cells^27,28^. To our knowledge, however, the role of ion exchange across gap junctions within multicellular systems has not yet been investigated. In this study, we develop a mechano-osmotic model to analyze the movement of fluid and ions between connected cells and their environment. Initially considering a simple two-cell system, we demonstrate that when a cell experiences increased solid stress loading, evolving osmotic pressure gradients drive swelling of its connected neighbor. We then expand our framework to explore how gap junctions amplify spatial variations in cell volume across a multicellular spheroid, highlighting an interplay between non-uniform proliferation-driven stress, cell mechanics, and transmembrane ion flow.

## Results

### A mechano-osmotic model for cellular volume control that integrates mechanical force balance with fluid and ion fluxes

To approach the problem of multicellular volume regulation, we initially consider two cells (Fig 1) held together via cadherin and catenin mediated complexes. As these complexes stabilize on the membrane, connexin structures also assemble and couple with identical units on the neighboring cell to form gap junctions. These channels connect the cytoplasm of both cells, permitting passive transport of fluid, ions, and small molecules^29^. Gap junctions typically remain open during their lifecycle, though may close in response to high Ca^2+^ concentrations or low pH which serves to protect the cell from dying neighbors^30^. Importantly, movement of fluid and ions not only depends on gap junction-mediated transport, but also on passive channels and active ion pumps within the cell membrane that permit transfer to and from the extracellular environment.

**Figure 1:**
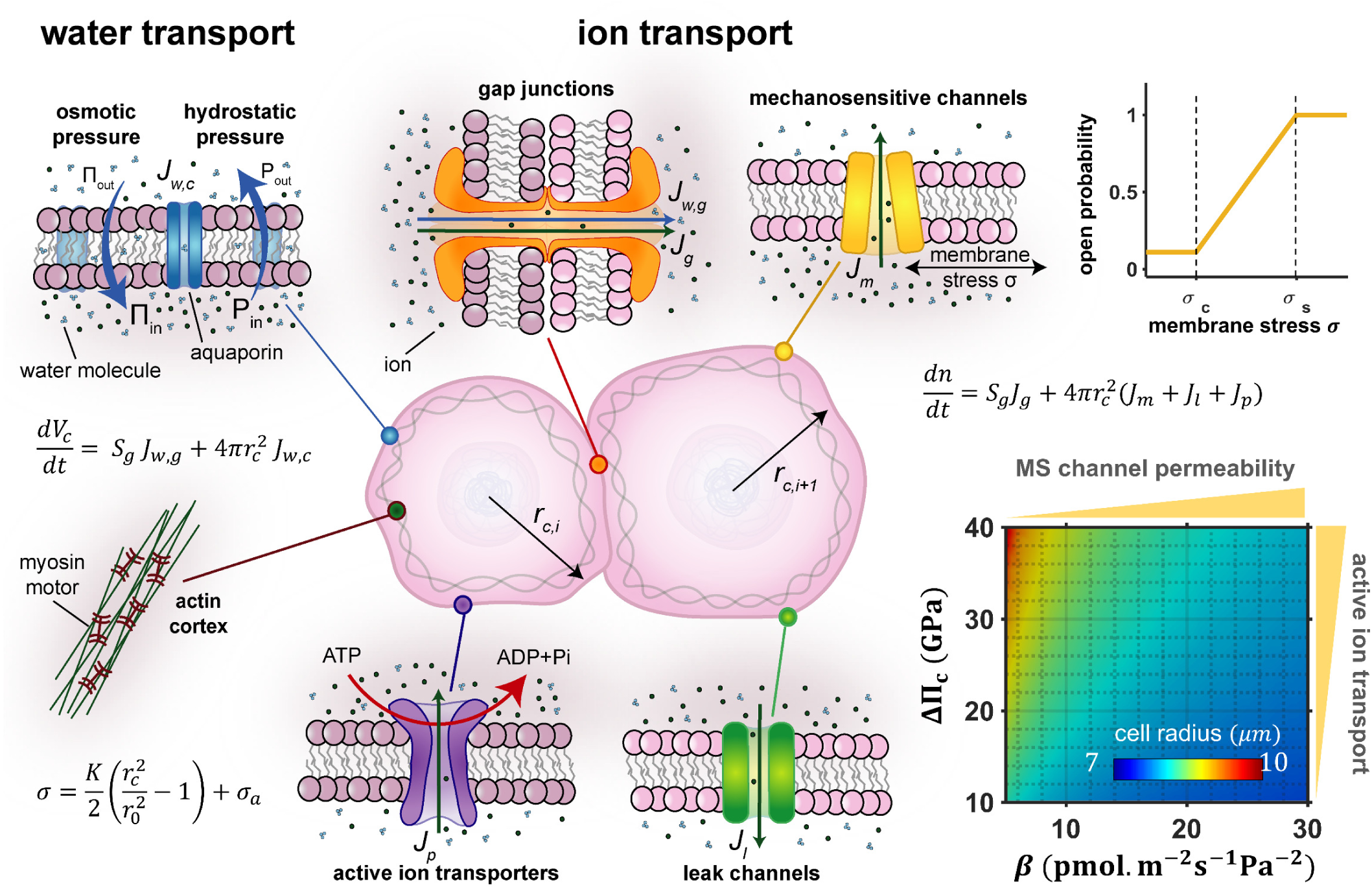
Schematic of cellular model. Water moves across the semi-permeable cell membrane as driven by hydrostatic or osmotic pressure gradients. Ions can diffuse between cells through gap junctions and across the membrane through mechanosensitive and leak channels. Active transporters move ions into the cell against a concentration gradient. Increased active transport or reduced channel permeability both lead to cell swelling.

### Gap Junction mediated ion and fluid transport between cells

To understand how water and ions move between the two cells, we first consider the role of hydrostatic and osmotic pressure, denoted by *P* and Π respectively. The cell’s internal osmotic pressure relates to the number of ions *n* in the cytosol via Van’t Hoffs equation Π = *nRT*/*V*_*c*_, where *R* is the gas constant, *T* is the absolute temperature, and *V*_*c*_ is the cell volume. In a cell *i*, the chemical potential of water depends both on the osmotic and hydrostatic pressure, given by Ψ_*w,i*_ = *P*_*i*_ − Π_*i*_. Movement of water between cells is driven by a difference in chemical potential ΔΨ_*w,g*_ across the gap junctions, such that the water flux may be described by *J*_*w,g,i*_ = −*ω*_*g*_ ΔΨ_*w,g*_ = −*ω*_*g*_(*P*_*i*_ − Π_*i*_ − (*P*_*i*+1_ − Π_*i*+1_)), where *ω*_*g*_ is a constant that relates to the water permeability of gap junctions. As cellular volume predominantly depends on its fluid content, we can therefore assume changes in volume are given by:

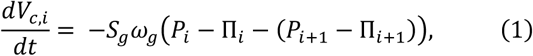

where *S*_*g*_ is the surface area of the membrane connected to the neighboring cell. Critically, in addition to water transport, gap junctions also permit diffusion of ions between cells. In the absence of external stimuli, Brownian motion of ions in fluid generates net diffusion into the cell with a lower ion concentration. This behavior may be characterized by a flux *J*_*g,i*_ = −*μ*(Π_i_ − Π_*i*+1_) where *μ* is a rate constant, and under these conditions the rate of change in the total number of ions in a cell can be determined:

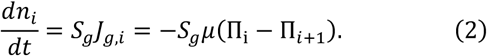

### Mechanics of the cell cortex

As water enters a cell, driven by hydrostatic and osmotic pressure gradients, the increase in fluid volume stretches the cell membrane. The mechanical tension in the membrane is complex, controlled by membrane-cytoskeleton adhesion, cortical stiffness, and active myosin contractility^31,32^. We treat the membrane and cortex as a single mechanical structure^26^, neglecting the possibility of cortical detachment and blebbing. The constitutive law of the cortical structure can be written as *σ*_*i*_ = *σ*_*p,i*_ + *σ*_*a,i*_, where *σ*_*a,i*_ is the active stress associated with myosin contractility and *σ*_*p,i*_ is the passive stress predominantly associated with deformation of the actin network (as the actin cortex is much stiffer than the plasma membrane^31,33^). With the assumption that the passive stress increases linearly with stretch, it can be expressed as 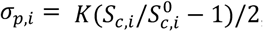, where *K* is the effective stiffness, *S*_*c,i*_ is the surface area of the cell, and 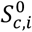 is a reference surface area. In addition to internal fluid pressure, the membrane also experiences loading from a spatially uniform external fluid pressure *P*^*ext*^. Mechanical force balance for a spherical cell with radius *r*_*c,i*_ dictates that the cortical stress can be related to the pressure difference across the membrane Δ*P*_*i*_ = *P*_*i*_ − *P*^*ext*^. Therefore, the cortical stress may also be written as *σ*_*i*_ = Δ*P*_*i*_*r*_*c,i*_/2*h*_*i*_, where *h*_*i*_ is the cortical thickness. Further, within a multicellular organoid, proliferation of cells generates compressive solid stresses *σ*_*g,i*_ that act on neighboring cells^34^. Deformation of fibrous matrix surrounding the cell cluster compounds the stress, as stretched fibers squeeze on the cluster^35^. Thus, we obtain the following expanded expression for the membrane/cortical stress:

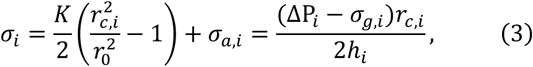

where *r*_0_ is the reference cell radius.

### Fluid and ion exchange with the extracellular environment

In addition to diffusion across gap junctions, water molecules can move through the semi-permeable cell membrane, enhanced by the presence of aquaporins^36^. This additional water flux across the membrane depends on the difference in osmotic and water pressure between the cell and the extracellular environment. We assume that the ion concentration in the external media is uniform, such that the external osmotic pressure at any point is given by Π^*ext*^. Therefore, the membrane water flux can be expressed as *J*_*w,c,i*_ = −*ω*_*c*_(Δ*P*_*i*_ − ΔΠ_*i*_), where Δ*P*_*i*_ = *P*_*i*_ − *P*^*ext*^ and ΔΠ_*i*_ = Π_*i*_ − Π^*ext*^, and *ω*_*c*_ is a rate constant^26^. We can extend Eqn 1 to consider this additional water flux such that *dV*_*c,i*_/*dt* = *S*_*g*_ *J*_*w,g,i*_ + *S*_*c,i*_ *J*_*w,c,i*_. Note that with our assumption of uniform external hydrostatic and osmotic pressures (e.g.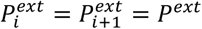) we can also state the gap junction water flux as a function of pressure differences, such that *J*_*w,g*_= Δ*P*_*i*_ − ΔΠ_*i*_ − (Δ*P*_*i*+1_ − ΔΠ_*i*+1_). Assuming the cells can be approximated to retain a spherical shape with radius *r*_*c,i*_, we achieve the following expanded form for cellular volume change:

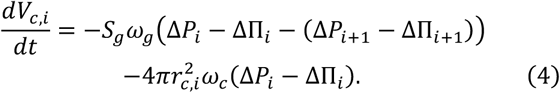

The cellular ion concentration also depends on exchange with the extracellular environment. Mechanosensitive (MS) channels are proteins in the cell membrane that open under a tensile membrane stress^37^ to allow flow of ions from regions where the concentration is high to regions where it is low. In response to hypotonic shock they release ions from the cell to mitigate an influx of water. The probability of channel opening has been reported to follow a Boltzmann function^38^, and consistent with previous work^26^, we adopt a piecewise linear expression (Fig 1, yellow curve) to describe the ion flux associated with MS channel permeability *J*_*m,i*_ = −*f*(*σ*_*i*_)ΔΠ_*i*_, such that

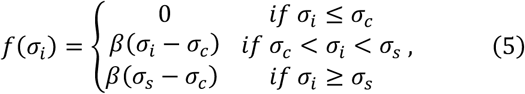

where *σ*_*c*_ is the threshold stress, below which *J*_*m,i*_ = 0, *σ*_*S*_ is the saturating stress, above which the channels are fully open, and *β* is a rate constant. In addition to these force sensitive channels, there are a number of leak channels (which are always operative) on the membrane^2^ for which we consider an further transmembrane ion flux *J*_*l,i*_ = −*ζ*ΔΠ_*i*_, where *ζ* is the associated rate constant. While the channels described thus far permit passive ion diffusion, there are additional membrane proteins present that actively transport ions against the concentration gradient. Such ion transporters require an energy input, such as from ATP hydrolysis, to overcome the energetic barrier associated with moving ions against the concentration gradient. We consider there to be a critical osmotic pressure difference the cell works to attain through active pumping^26^, given by ΔΠ_*c*_ = Π^*ext*^ Δ*G*_*a*_/*RT*, where Δ*G*_*a*_ is the required energy input. Thus the ion flux generated by active pumping by ion transporters can be expressed as *J*_*p,i*_ = *γ*(ΔΠ_*c*_ − ΔΠ_*i*_), where *γ* is a rate constant. Taking these pumps and channels into consideration, we can extend Eqn 2 for a more detailed description of the number of ions within the cell whereby *dn*_*i*_/*dt* = *S*_*g*_*J*_*g,i*_ + *S*_*c,i*_(*J*_*m,i*_ + *J*_*l,i*_ + *J*_*p,i*_), such that

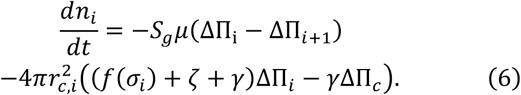

Increasing active ion transport or decreasing MS channel permeability will increase the cytosolic ion concentration, leading to an increase in cell volume (Fig 1). Material parameters for all simulations are summarized in Table S1.

### Gap junctions amplify differential swelling associated with proliferation-induced solid stresses in neighboring cells

Typically, cell clusters are seeded in a confining matrix, and as the cluster grows it displaces and deforms the elastic matrix, opposing growth and generating solid stresses within the cluster. When cells proliferate within the growing cluster, they push against and apply compressive stresses on their neighbors^34^, and with the addition of cell adhesion and local jamming this leads to different levels of stress developing spatially. To understand the influence of such solid growth stress on cellular shrinkage and swelling, we first consider the interactions between two connected cells (Fig 2A). Without loss of generality, we assume there to be a compressive stress *σ*_*g*,1_ = *σ*_*g*,0_ + *δσ*_*g*,1_ acting on one cell and for its neighbor the be acted upon by a stress *σ*_*g*,2_ = *σ*_*g*,0_. For the purposes of illustration, we choose *σ*_*g*,0_ to equal zero and for *δσ*_*g*,1_ to increase over time to a maximum of 50 Pa (Fig S1A). Varying these parameters will lead to similar trends albeit different magnitudes.

**Figure 2:**
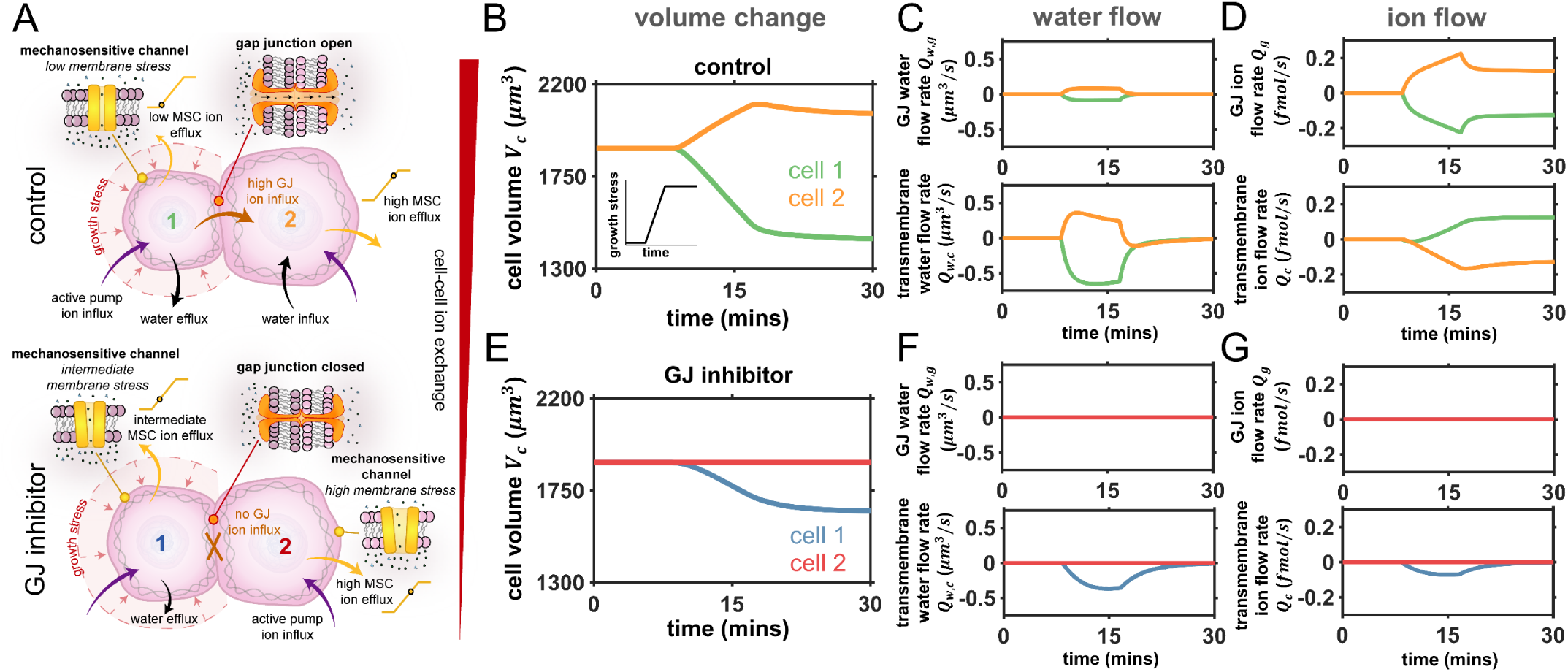
Role of gap junctions (GJ) in cellular volume control. A) Schematic of solid growth stress driving swelling in neighboring cell which is prevented by inhibition of ion transport across GJs; B) In the control case the loaded cell shrinks and the neighboring cell swell. Inset shows the loading profile that increases over time; C) Control case water flow rate *Q*_*w*_ = *S J*_*w*_ across gap junctions and the membrane; D) Control case ion flow rate *Q* = *S J* across gap junctions and the membrane; E) When ion transport across gap junctions is inhibited (*μ* = *ω*_*g*_ = 0) there is negligible volume change in the neighboring cell; F) Inhibition case water flow rate *Q*_*w*_ = *S J*_*w*_ across gap junctions and the membrane; G) Inhibition case ion flow rate *Q* = *S J* across gap junctions and the membrane.

When gap junctions are active (control case), our model predicts that the loaded cell shrinks and its neighbor swells (Fig 2B). Initially, in the absence of loading, we find that cells control their volume by regulating their ion concentration through the activities of ion pumps and channels (Fig 2B; Fig S1). In this unloaded state, MS channels are permeable due to tension in the cell membrane, permitting a constant loss of ions to the external media (Fig S1D). However, active ion pumping ensures there is a continuous ion influx (Fig S1F) to maintain the cell’s osmotic pressure higher than that of the external media, allowing it to retain water. When solid stress on the cell’s surface increases, its internal hydrostatic pressure also increases (Fig S1B) and water is squeezed from the cell (Fig 2C); recall that water flow is partly driven hydrostatic pressure differences (Eqn 4). The loss of water relieves tension in the cell’s membrane, thereby reducing the permeability of its MS channels (via Eqn 5). However, as ion pumping remains active there is a continuous intake of ions (Fig S1F). Overall, the cell loses fewer ions, but the rate of ions entering the cell remains relatively constant and therefore there is a net increase in its ion concentration. This increases the loaded cell’s osmotic pressure (Fig S1C) relative to its connected neighbor, and the difference generates a flow of ions through their gap junctions from the loaded to the unloaded cell (Fig 2D). The resulting increase in the unloaded cell’s ion concentration causes it to absorb additional water from the external media and swell (Fig 2B). Although its MS channels are now more permeable due to increased membrane tension, the constant flow of ions from the loaded cell maintains the neighbors swollen state. Thus, compression of a cell increases its osmotic pressure relative to its connected neighbor due to closing of mechanosensitive channels. This drives a flow of ions across gap junctions into the unloaded cell, causing it to absorb water and swell.

To further analyze the role of cell-cell transport in volume regulation, we inhibit gap junctions by reducing their ion and water permeability (via *μ* and *ω*_*g*_, respectively) to zero. The loaded cell shrinks (Fig 2E), but it loses less water than it would under control conditions. This reduction in volume again relieves membrane tension, reducing the permeability of MS channels and increasing the cell’s overall ion concentration. However, as gap junctions are blocked there is no loss of ions to the neighboring cell (Fig 2G) and thus the loaded cell sustains its high osmotic pressure (Fig S2C). This allows it to retain more water (and maintain a high hydrostatic pressure) to oppose the applied compressive stress. Additionally, there is no swelling predicted in the neighboring cell (Fig 2E), owing to the lack of cell-cell ion exchange. Clearly, ion transport across cellular gap junctions (GJ) plays an important role in cellular volume regulation, with an increasing GJ permeability driving larger differences in cell volume (Fig S3).

To assess if cell swelling could be driven by intercellular fluid flow alone (i.e. no ion transport) we also investigate inhibition of the GJ ion flux while maintaining intercellular movement of water (Fig S4). Fluid is initially squeezed into the neighboring cell in response to loading (Fig S4C). However, the additional water intake stretches the membrane of the neighboring cell, increasing its internal hydrostatic pressure and driving the fluid into the external media (Fig S4D). Thus, our model suggests that intercellular water flow alone does not drive significant cell swelling.

### Interstitial fluid pressure has a secondary influence on volume dynamics in connected cells

Importantly, in our analyses we have implicitly assumed that the extracellular hydrostatic fluid pressure *P*^*ext*^ is spatially uniform (i.e.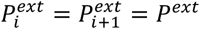) and that cell loading is predominantly attributed to solid stress. However, within proliferating tumors there may also be local interstitial pressure gradients that can influence cell behavior. Thus, here we explore their influence on cell shrinkage and swelling. Recall that the water flux across the cell membrane is driven by a balance between the internal and external hydrostatic and osmotic pressures, such that *J*_*w,c,i*_ = −*ω*_*c*_(Δ*P*_*i*_ − ΔΠ_*i*_), where Δ*P*_*i*_ = *P*_*i*_ − *P*^*ext*^ and ΔΠ_*i*_ = Π_*i*_ − Π^*ext*^. Without loss of generality this flux can be rephrased such that 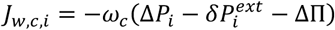, where 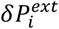 is a hydrostatic pressure perturbation acting only on cell *i*. Thus, extending Eqn 4, the change in cell volume may be expressed as:

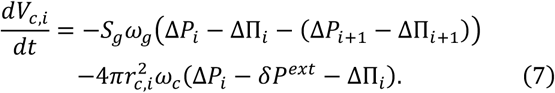

As an increase in the interstitial fluid pressure will also affect loading on the cell membrane, the mechanical force balance must also be updated:

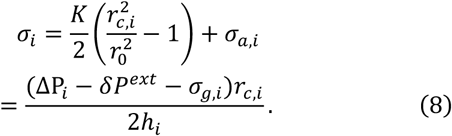

In the absence of gap junctions, our model predicts that (unlike solid stress) an additional fluid pressure (Fig S5A) does not cause a change in cell volume (Fig S5B-solid line). Although the increase in external hydrostatic pressure should drive water into the cell, swelling is opposed by the compressive load that the same interstitial pressure imposes on the cell membrane. However, the internal hydrostatic pressure of the loaded cell increases (by a magnitude equal to *δP*^*ext*^ (Fig S5C)). When GJs become permeable, because the hydrostatic pressure of the loaded cell is higher than that of its neighbor, water flows into the neighboring cell (Fig S5E). The loaded cell then shrinks due to water loss and its MS channels become less permeable, limiting ion efflux and increasing osmotic pressure. Similar to the mechanism outlined for solid stress loading, this then generates a flow of ions across the gap junctions, increasing the ion concentration in the neighboring cell and causing it to swell. The volume change is predicted to become increasingly pronounced with increasing GJ permeability. In summary, with active gap junctions solid stress and interstitial fluid pressure are both predicted to cause shrinkage of the loaded cell and swelling of the connected neighbor. However, it is important to note that under identical conditions the magnitude of the volume change is significantly (∼10-fold) higher in response to solid stress (Fig S6). In the next section, we proceed to generalize our analysis of volume changes in a two-cell system to a multicellular cluster.

### Intercellular ion transport drives spatial variations in cell volume within proliferating solid tumors

Multicellular spheroids are an increasingly promising experimental model for cancer development, that aim to bridge the gap between in-vitro and in-vivo conditions^10^. In such 3D environments however, it remains unclear how individual cells regulate their volumes and coordinate to advance tumor progression and matrix invasion. Therefore, we next consider how our mathematical framework can be extended to predict fluid and ion exchange within connected cells in a tumor organoid (as driven by differences in solid stress, hydrostatic pressure, and osmotic pressure). While our current model can readily be adapted to simulate a series of discrete connected cells, more physical insights can be gained from a continuum formulation that describes cellular behavior within a spherical organoid (Fig 3A). Continuity requires that for any cell within a cluster, the change in its number of ions must equal the amount gained and lost through gap junctions to neighbors and through the cell membrane. First, considering cell-cell ion exchange, recall that the ion flux across gap junctions between two connected cells depends on their osmotic pressure difference, with *J*_*g,i*_ = −*μ*(ΔΠ_i_ – ΔΠ_*i*+1_). The gradient of osmotic pressure between these cells may be written as ∇(ΔΠ) = (ΔΠ_*i*_ − ΔΠ_*i*+1_)/*r*_0_. For a cell within a longer series, this can be extended to develop an expression for the Laplacian of the osmotic pressure, whereby:

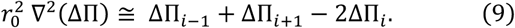

**Figure 3:**
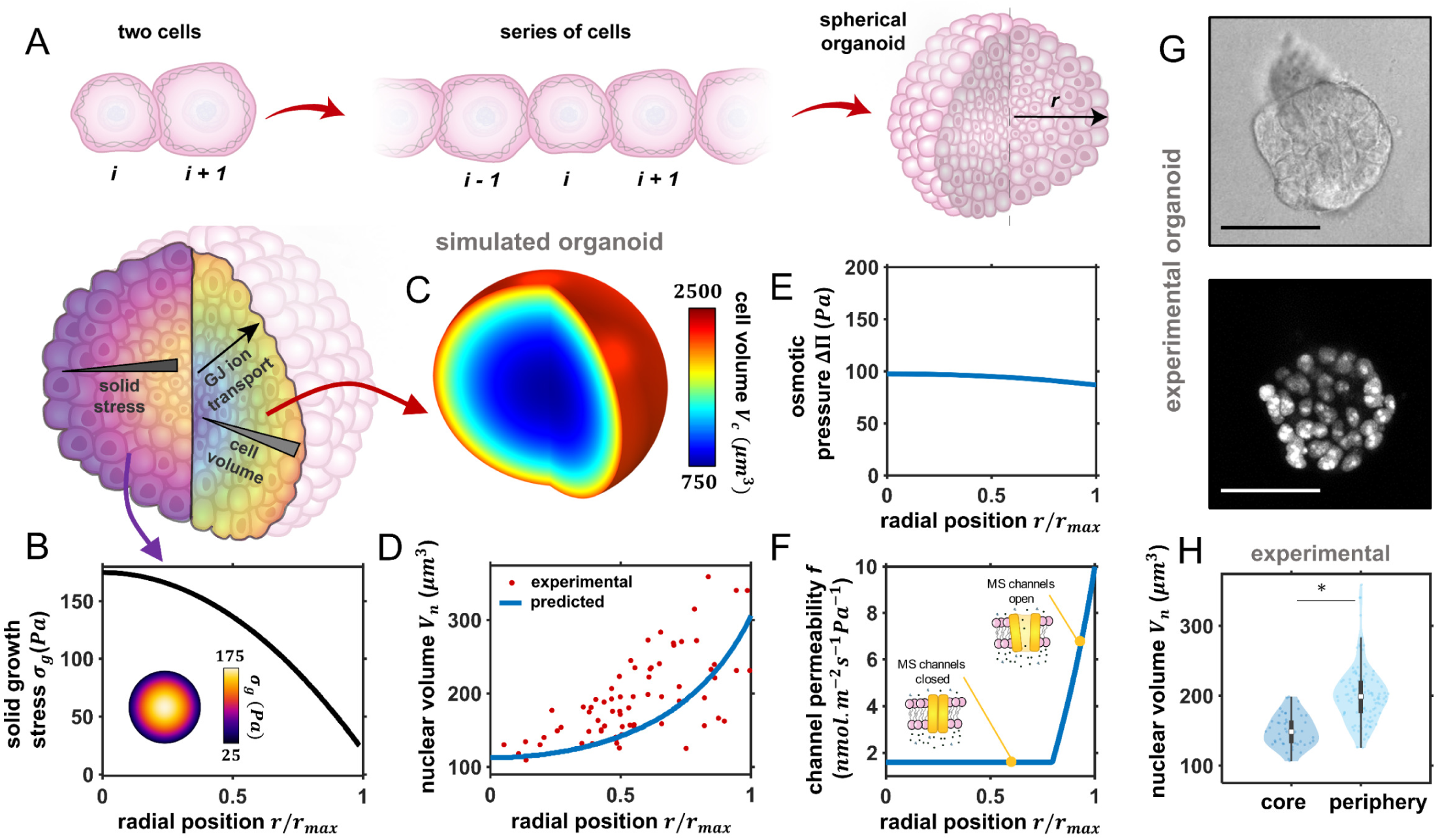
Spatially non-uniform cell volume in cancer organoid. A) Schematic of model extension to 3D spherical organoid; B) Applied solid growth stress *σ*_*g*_(*r*) is highest at the core and spatially non-uniform; C) Predicted spatial cell volume; D) Predicted and experimental (day 5) spatial nuclear volume; Predicted spatial E) osmotic pressure ΔП and F) channel permeability *f*′ = *f*(*σ*) + *ζ*; G) Cross-section images of epithelial cancer organoid developed from MCF 10A cells at day 5 of growth with GFP-NLS-labelled cell nuclei. Scale bar 50 *μm*; H) Nuclear volume of cells at organoid core and periphery at day 5 of growth. The boxes represent the interquartile range between the first and third quartiles, whereas the whiskers represent the 95% and 5% values and the squares represent the median. \**P* < 0. 001.

In a spherical coordinate system, assuming circumferential symmetry, this Laplacian can be rephrased as 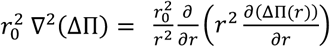, where *r* is the radial position in a spherical organoid. Further, the ion flux across gap junctions may also be extended to describe a cell in-series and connected on both sides, such that *J*_*g,i*_ = −*μ*((ΔΠ_i_ − ΔΠ_*i*-1_) + (ΔΠ_i_ − ΔΠ_*i*+1_)). Altogether, we can then enforce continuity to formulate a continuum expression that describes the number of ions entering and leaving a cell at position *r* within multicellular spheroid, such that:

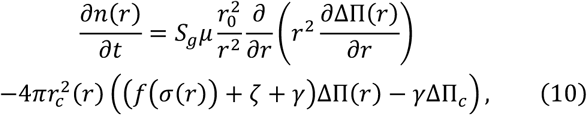

where the first term on the right describes the ions gained and lost through gap junctions in a cell at position *r*, and the second term accounts for cellular exchange of ions with the surrounding media via pumps and channels. Similarly, cellular volume change may be expressed as:

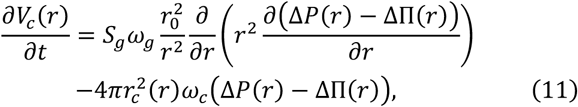

where, for a cell at position *r* in the organoid, the first term on the right accounts for the flow of water through gap junctions, and the second term describes water entering and leaving the cell through its membrane. We solve our system of equations at steady state (i.e.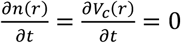) using multi-physics software COMSOL to simulate local cell behavior within a spherical organoid of radius *r*_*max*_ (see Methods for more details). As the gap junction flux vanishes at the cluster boundary, a zero-flux condition is enforced on the spheroid surface.

Proliferation of cells within a growing cluster generates solid compressive stresses, additionally compounded by matrix fiber tension and cell confinement. Interestingly, it has been shown that such stress is spatially non-uniform across the cancerous structure^39,40^, frequently highest at the cluster core and lowest at the periphery, with maximum stress 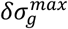 typically lying in the range of 100 − 250 *Pa*. Thus, we apply a spatially varying solid growth stress to the organoid, as shown in Fig. 3B, whereby the stress is highest at the core and decreases radially such that 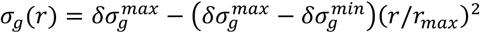; alternative stress distributions are explored in Fig S7. As we have identified that a spatially varying interstitial fluid pressure has a significantly lower influence on volume changes than solid stress (Fig S6), we approximate the external hydrostatic pressure to be spatially uniform.

Our model predicts that cell volume is lowest at the core (*V*_*c*_(*r* = 0) = 750 *μm*^3^) and increases radially (*V*_*c*_(*r* = *r*_*max*_) = 2500 *μm*^3^), as shown in the organoid contour plot in Fig. 3C. We have shown that the ratio of nuclear to cell volume remains constant during volume changes^11^, maintaining a value of *V*_*n*_: *V*_*c*_ ≅ 0.14. Applying this ratio, we can predict nuclear volumes across the organoid (Fig 3D), and show strong agreement with our recent experiments^11^ as discussed in more detail in the next section. At the core, where there is the most significant cell shrinkage, water loss reduces the membrane tension and thus MS channel permeability is impaired (Fig 3F). As a result, the osmotic pressure of internal cells increases (Fig 3E). Radially, the solid growth stress reduces, and ion channels become increasingly permeable provided the membrane stress exceeds the critical value *σ*_*c*_ (Eqn 5). Within the whole organoid, this generates an intercellular ion concentration gradient that propagates radially (Fig 3E), driving an ion flow that increases the osmotic pressure in peripheral cells and causes them to absorb water. The loss of ions from core cells impedes their ability to retain water, and thus they are highly compressed during loading. As such, at steady state there is a balance attained between the solid growth stress (that drives a water efflux) and cellular ion concentrations (that drive a water influx). During an early stage of organoid growth when solid stress is low and approximately uniform, there is no predicted spatial variation in cell volume (Fig S8). In summary, our model suggests that ion flow through gap junctions, as generated by differences in cellular ion concentrations, is a significant driving factor in cell swelling and shrinkage within multicellular organoids. In the next section, we discuss our recent experiments to validate these predictions.

### Experimental evidence validates model predictions that gap junctions mediate cell swelling in multicellular spheroids

In recent work, we experimentally uncovered that locality within a cancer organoid governs cell volume and stiffness^11^. We seeded single MCF 10A human breast epithelial cells into 3D hydrogels composed of 4 *mg ml*^-1^ Matrigel and 5 *mg ml*^-1^ alginate (see Methods for details), such that the gels had a shear modulus of approximately 300 *Pa* to reflect the environment of in-vivo breast carcinoma. Initially, an isolated cell proliferated to form a spherical cluster (day 3), continuing to grow into a larger spheroid (day 5) with cells present both in the core and at the periphery (Fig 3G).

Further growth led to invasive branches extending into the surrounding matrix. Cells were transfected with a green fluorescent protein (GFP) tagged nuclear-localization signal (NLS), and also labelled using cytoplasmic staining, which enabled measurement of cell and nuclear volume using 3D confocal microscopy (Fig 3G). Importantly, we identified that the ratio of nucleus to cell volume remained constant over a wide range of organoid sizes and cell positions, in agreement with previous findings^41,42^, which allowed us to measure nuclear volume in lieu of cell volume. At an early stage of growth, all cells had similar nuclear volumes (Fig S8A). However, as the organoid further developed, nuclear volume began to correlate strongly with cell position within the cluster. Cells towards the core (inner 40% of organoid radius) were significantly smaller than those in peripheral regions (Fig 3H), with a volume range in strong agreement with our model predictions (Fig 3D). Interestingly, we also demonstrated that a reduction in volume was highly correlated with an increase in cell stiffness (measured using optical tweezers active microrheology^43^), most likely due to increased molecular crowding^41,44^. These experimental results both validate our model findings and highlight the importance of understanding the mechanisms of volume change within multicellular spheroids.

As our model suggests that GJs are a critical mediator of spatial volume variation within cancer organoids, we next explore how cell behavior changes when GJs are blocked. The spatial variation in cell volume is shown to decrease significantly (Fig 4A), ranging from ∼1500 *μm*^3^ at the core to ∼1700 *μm*^3^ at the periphery. As per the control (active GJs) case, compressive stress reduces cell volume throughout the spheroid. However, when the MS channels of inner cells close (Fig 4D), there is no loss of ions through GJs to relieve the cells’ high ion concentrations. Thus, relative to the control case, cellular osmotic and hydrostatic pressure is significantly higher at the organoid core (Fig 4E). Further, peripheral cells have a low osmotic pressure because they do not gain ions from their neighbors; therefore they do not absorb water and swell. In fact, they retain a volume similar to that during an early stage of organoid growth (Fig S8A). In our experiments, we inhibited GJs by adding 500*μM* carbenoxolone to the organoid-matrix system on day 3 (before a volume gradient was present)^11^. In agreement with our simulations, on day 5 we did not observe significant spatial differences in nuclear volume (Fig 4A,C). It is important to note that intercellular water flow through GJs is not sufficient to drive the changes in cell volume, as the additional fluid would simply diffuse through the cell membrane in response to hydrostatic pressure gradients (Fig S4). This further supports our model findings that ion diffusion driven by an intercellular osmotic pressure gradient is the critical factor in cell swelling and shrinkage within the organoid.

**Figure 4:**
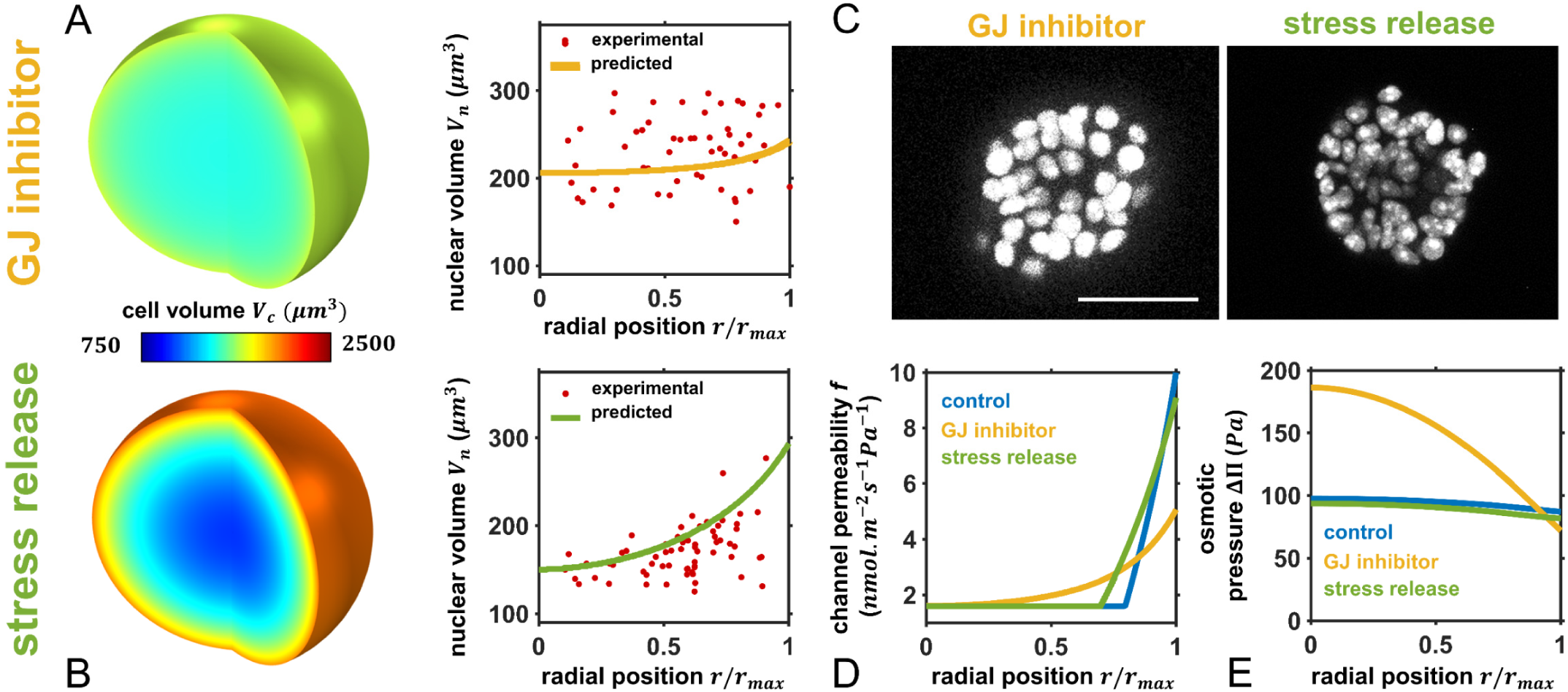
Influence of GJ inhibition and stress release on cellular swelling. Predicted and experimental (day 5) spatial cell and nuclear volumes under A) inhibited gap junction and B) stress release conditions; C) Cross-section images of GFP-NLS-labelled MCF 10A cells at day 5. Scale bar 50 *μm*; Predicted spatial D) channel permeability *f*′ = *f*(*σ*) + *ζ* and E) osmotic pressure ΔΠ.

Finally, as cell volume clearly additionally depends on the solid stress gradient, we examined the influence of reducing the solid stress acting on the cell cluster. Our model predicts that a reduction in the maximum (core) solid stress (profile shown in Fig S9A) also reduces the spatial variation in cell volume (Fig 4B). The reduction in stress allows more cells to sustain their MS channel permeability (Fig 4D), permitting a loss of ions to the interstitium and thereby lowering their osmotic pressure. This reduces intercellular ion diffusion and thus lowers cell swelling in peripheral regions of the cluster. We performed additional experiments that can test these predictions, whereby using organoids cultured in a collagen (3.5 *mg ml*^-1^) / Matrigel (0.5 *mg ml*^-1^) matrix, we reduced solid stress on day 5 by degrading collagen fibers with collagenase. After 6 hours, we observed a significant reduction in nuclear volume at the periphery and an increase at the core (Fig 4B,C); the volume gradient became weaker, in agreement with model findings. Therefore, the spatial variation in cell volume within the cancer organoid may be understood to depend on intercellular ion diffusion in response to an osmotic pressure gradient, as driven by non-uniform external solid stress.

## Discussion

In this study, we propose a mechano-osmotic model to investigate how cell volume is regulated within multicellular systems. Volume control depends on an interplay between multiple cellular constituents, including gap junctions, mechanosensitive ion channels, energy consuming ion transporters, and the actomyosin cortex, that coordinate to manipulate cellular osmolarity. In connected cells, mechanical loading is shown to significantly affect how these components cooperate to transport ions, and precise volume control is impacted by the evolution of osmotic pressure gradients between cells. Combining the modeling framework with our recent experiments, we ultimately identify how gap junctions can amplify spatial variations in cell volume within multicellular spheroids and, further, describe how the process depends on proliferation-induced solid stress (Fig 5). Initially considering a simple two-cell system, our model predicts that compressive stresses squeeze water from a cell, and a subsequent reduction in membrane tension and loss of mechanosensitive channel permeability impedes the flow of ions to the external media. As ion transporters continue to pump new ions into the cell, there is an effectual increase in the cytosolic ion concentration. In connected cells, a flow of ions through gap junctions is thus generated from the loaded cell, thereby increasing the osmolarity of its neighbor and causing it to swell. Blocking of gap junctions is revealed to prevent this volume change and, also, reduce shrinkage of the loaded cell.

**Figure 5:**
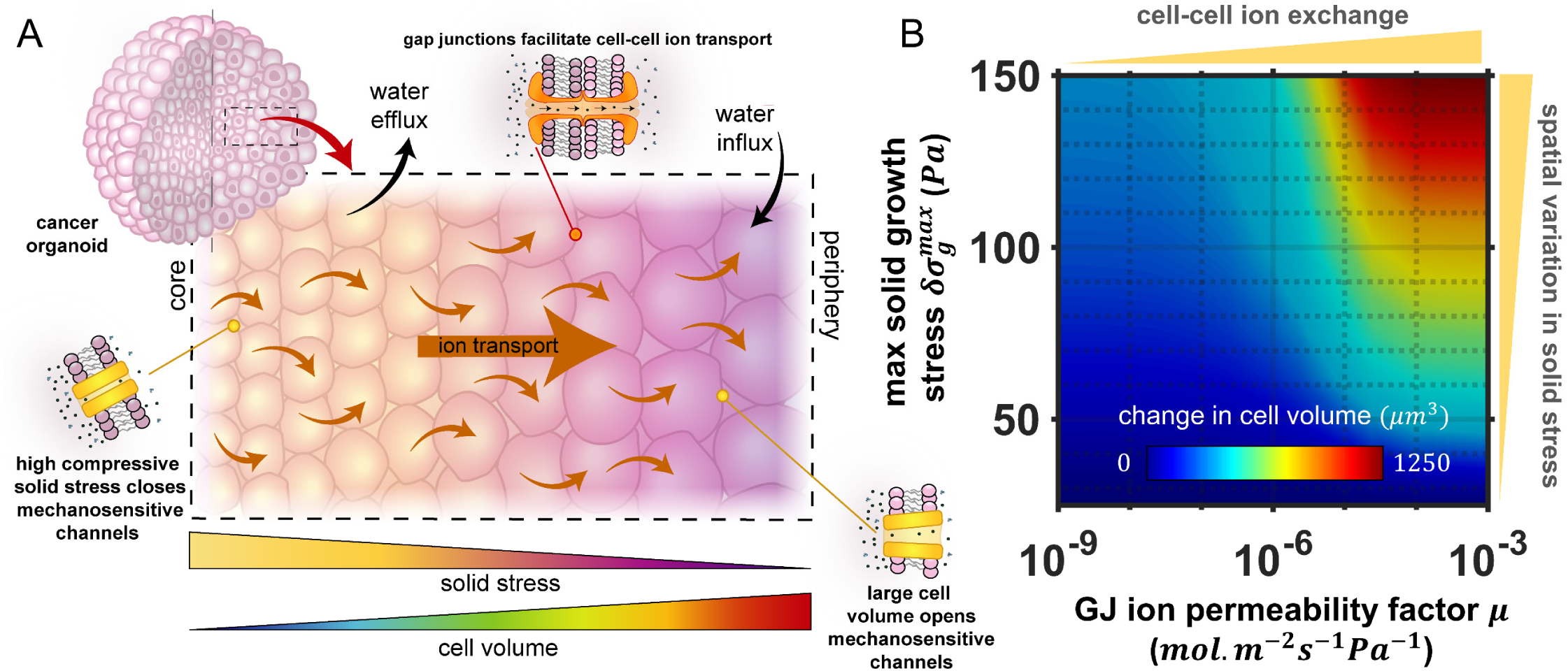
A) Non-uniform solid growth stress drives spatial variation in cell volume via gap junction mediated ion flow; B) Spatial variation in cell volume is increased both by high cell-cell ion transport and high solid growth stress.

We next extended our framework to explore how spatial variations in cell volume can emerge within a multicellular tumor. Cancer cells are typically situated within a confining matrix, and as the cells proliferate they both push against their neighbors and displace the elastic matrix. Associated solid stresses develop within the cluster, generally evolving to be highest at the cluster core^39,40^. In response, core cells tend to be significantly compressed and lose volume. This relieves their cellular membrane tension, which closes mechanosensitive ion channels, and a reduction in this channel permeability dictates that fewer ions are lost to the interstitium. Overall, as ions are still pumped into the cells via active transport, there is a net increase in their internal ion concentration. With a radial reduction in solid stress loading, more ion channels remain open towards the organoid periphery and thus peripheral cells maintain lower osmotic pressures. Following the mechanisms uncovered from our two-cell analysis, a radial intercellular flow of ions is generated from cells in the core, thereby increasing the ion concentration in cells situated towards the periphery. These cells then swell in response to a water influx driven by their increased osmolarity (Fig 5A). Interestingly, from our continuum expressions (Eqn 10-11) we can identify an effective length scale for cell volume changes in response to solid stress 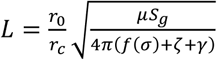, which reveals that the transmission distance increases with either increasing GJ permeability *μ* or reduced MS channel permeability *f*(*σ*). Further, we can readily obtain analytical solutions for our continuum formulations at the limits of GJ permeability (SI Note 1), to highlight that GJs can amplify spatial differences in cell volume. In fact, our simulations clearly show that increasing GJ permeability promotes larger volume differences across the multicellular spheroid (Fig 5B). Spatial variation in cell volume is also amplified at higher solid stress gradients.

Gap junctions play a critical role in supporting many physiological operations, including embryonic development^45^ and collective cell migration^46^. Clearly, our framework could be used to analyze such biological systems and provide mechanistic insight into their dependence on ion transport and differential cell swelling. Future advancements should also focus on detailing the interdependence between dynamic actomyosin contractility and cell osmolarity, building on our previous work to further understand how the two-way feedback between stress and signaling guides nuclear gene expression^47^, dynamic force generation^48^, and cancer invasion^13^. Importantly, in the current analysis we limit ourselves to the consideration of a single ion species and the associated channels and transporters, and therefore neglect the requirement of charge neutrality. This simplification of the complex cellular transport system allowed us to identify the critical processes that can drive spatial variations in volume within proliferating cell clusters. Nonetheless, our framework can readily be extended to consider m ultiple solute species and the influence of voltage-gating on ion transport to explore more specific solute and protein interactions.

In cancer progression, the precise influence of gap junctions remains a point of avid debate^49^. In early studies, loss of intercellular communication was identified to be characteristic of cancer cells ^50^. However, it has been reported that low expression of the GJ protein connexin correlates with both positive and negative prognoses across a range of cancer types^51,52^. Particularly in late stage tumors, there is an increasing body of evidence that indicates expression of connexin is associated with tumor malignancy, growth, and invasion^53–55^. In this study, we identified that ion flow through gap junctions promotes peripheral cell swelling in loaded tumor organoids. As controlled swelling has been shown to increase cell proliferation^18,19^, this could hint at a mechanism by which GJs support cancer growth. Further, in our recent experiments we demonstrated that swelling correlated with reduced cell stiffness^11^, which has been suggested as a metastatic biomarker associated with increased invasive potential^56^. Accordingly, we found that osmotic swelling increased cell invasiveness, while blocking GJs led to a reduction in the number of invading branches, indicating that GJ mediated swelling promotes matrix invasion. In summary, our findings suggest that intercellular ion flow may be an important mediator of cancer progression. Our proposed model can help guide the development of therapeutics that target inter- and extra-cellular ion transport.

## Methods

### Simulation procedure

For the two-cell dynamic analysis the model was implemented using an ODE solver (ode23s) in MATLAB (v. 2019a, MathWorks). Initial conditions were identified by solving Eqns 1-6 at steady state. Cell behavior was simulated over 30 minutes, with a time-dependent load *σ*_*g*,1_ introduced following 8 minutes (Fig S1A). All material parameters are summarized in Table S1. For the cell cluster analysis a 2D axisymmetric spheroid model of radius *r*_*max*_ was constructed in multi-physics software COMSOL (v. 5.4, COMSOL AB). Using in-built PDE solver functionality Eqns 10-11 were solved in conjunction with mechanical equilibrium (Eqn 3) to determine spatial steady state cell behavior in response to an applied non-uniform load *σ*_*g*_(*r*). A zero-flux condition was enforced on the spheroid surface.

### Cell lines and cell culture

MCF10A cells (ATCC, CRL-10317) were cultured in complete medium at 37 °C with 5% CO_2_. The complete medium is made of DMEM/F12 medium (Invitrogen, 11965-118) supplemented with 5% horse serum (Invitrogen, 16050-122), 20 ng/ml epidermal growth factor (Peprotech, AF-100-15), 0.5 μg/ml hydrocortisone (Sigma, H-0888), 100 ng/ml cholera toxin (Sigma, C-8052), 10 μg/ml insulin (Sigma, I-1882) and 1% penicillin and streptomycin (Thermo Fisher, 15140122). We transfected the MCF10A cell line with GFP-NLS lentivirus to visualize the cell nucleus following the product manual (Essen Bioscience, 4475), and the stable cell line was maintained in T-25 cell culture flask with complete medium and 0.4 mg/mL puromycin (Thermo Fisher, A1113802). Subculture was performed when cells grow into 80% confluency. Briefly, cells were washed with PBS for three times before 1 mL of 0.05% trypsin-EDTA solution (Thermo Fisher, 25300054) was added. Then the T-25 flask was incubated at 37 °C for 15 min. After most of the cells detached from the flask, cells were collected, centrifuged (1500 rpm, 5 min) and resuspended into a new flask.

### Growth of MCF10A clusters

The MCF10A clusters with invasive phenotype were cultured and induced following previously established protocols^57,58^. Briefly, two hydrogel systems were used in this study for 3D cell culture, including alginate-Matrigel hydrogel and collagen-Matrigel hydrogel. For alginate-Matrigel hydrogel, cells were mixed with alginate (FMC Biopolymer), calcium sulfate (Sigma, 255696) and Matrigel (Corning, 354234) to form gel precursor solution with final concentrations of 5 mg/ml, 20 mM and 4 mg/ml, respectively. For collagen-Matrigel hydrogel, cells were mixed collagen (Advanced BioMatrix, 5133) and Matrigel with final concentrations of 3.5 mg/ml, 0.5 mg/ml, respectively. The gel precursor solution was then incubated in 37 °C for 30 min to form cell-laden hydrogel and cultured in complete cell culture medium for 10 day. To inhibit gap junctions, 500 µM carbenoxolone was added to the complete cell culture medium. To reduce solid stress within the clusters in collagen-Matrigel system, collagenase D (Sigma, 11088866001) was used to remove the matrix.

### Nuclear volume measurements

The 3D structure of the MCF10A clusters was imaged with a confocal microscopy (Leica, TCL SP8), and deconvolution (HUYGENS software) was applied to the image to improve the z resolution of traditional confocal microscopy. The nuclear volume was then calculated by the number of voxels contained within the nuclear structures using a customized algorithm in MATLAB (v. 2017a, Mathworks).

### Statistics

A two-tailed Student’s t-test was used when comparing the difference between two groups. In the box plots, the boxes represent the interquartile range between the first and third quartiles, whereas the whiskers represent the 95% and 5% values, and the squares represent the median.

## Data Availability

Data supporting the findings of this study are available within the article and the Supplementary Information, and source data are available from the corresponding author on reasonable request.

## Code Availability

MATLAB and COMSOL files used in this work are available from the corresponding author on reasonable request.

## Acknowledgements

This work was supported by National Cancer Institute Awards U01CA202177 (V.B.S.), U54CA193417 (V.B.S.), R01CA232256 (V.B.S) and U01CA202123 (M.G.); National Institute of Biomedical Imaging and Bioengineering Award R01EB017753 (V.B.S.); NSF Center for Engineering Mechanobiology Grant CMMI-154857 (V.B.S.); NSF Grant MRSEC/DMR-1720530 (V.B.S.); Alfred Sloan Research Fellowship (M.G.).

## Author contributions

E.Mc. and V.B.S. designed the theoretical models and carried out the computations; Y.H and M.G. designed and conducted the experiments; E.Mc., Y.H., M.G., and V.B.S. analyzed and interpreted the data; E.Mc., Y.H., M.G., and V.B.S. prepared and edited the manuscript.

