## Supplementary for "Gap Junctions Amplify Spatial Variations in Cell Volume in Proliferating Solid Tumors"

| Parameter | Description | Value |
|---|---|---|
| $\mu$ | gap junction ion permeability factor ($mol. m^{-2}s^{-1}Pa^{-1}$) | $5\ X\ 10^{-7}$ |
| $\omega_g$ | gap junction water permeability factor ($m. s^{-1}Pa^{-1}$) | $10^{-9}$ |
| $r_0$ | reference cell radius ($\mu m$) | 7.1 |
| $S_g$ | area of membrane adhered between two cells ($\mu m^2$) | 63.3 |
| $K$ | effective stiffness of cortical layer ($kPa$) | $3^*$ |
| $\sigma_a$ | active cortical stress ($Pa$) | $100^*$ |
| $h$ | thickness of cortical layer ($\mu m$) | $0.5^*$ |
| $\beta$ | MS channel ion permeability factor ($mol. m^{-2}s^{-1}Pa^{-2}$) | $2\ X\ 10^{-11*}$ |
| $\sigma_c$ | threshold stress of MS channel ($Pa$) | 75 |
| $\sigma_s$ | saturating stress of MS channel ($Pa$) | 600 |
| $\zeta$ | leak channel ion permeability factor ($mol. m^{-2}s^{-1}Pa^{-2}$) | $1.5\ X\ 10^{-9}$ |
| $\gamma$ | rate constant of ion flux via active transporters ($mol. m^{-2}s^{-1}Pa^{-1}$) | $10^{-17*}$ |
| $\Delta\Pi_c$ | critical osmotic pressure difference of ion pump ($GPa$) | $30^*$ |
| $\Pi^{\text{ext}}$ | external osmotic pressure ($MPa$) | $0.5^*$ |
| $\omega_c$ | cell membrane water permeability factor ($m. s^{-1}Pa^{-1}$) | $10^{-9*}$ |
| $r_{max}$ | spheroid radius ($\mu m$) | 33.4 |

**Table S1: *Parameters for chemo-osmotic model.* The reference cell radius $r_0$ was approximated such that the prediced cell volumes at an early stage of organoid growth provide good agreement with our experiments (Fig S8). The spheroid radius $r_{max}$ was directly measured from the day 5 experimental images (Fig 3). The gap junction ion permeability factor $\mu$ was estimated assuming that the ion flow rate across the membrane and gap junctions operated within an order of magnitude (Fig 2), and the gap junction water permeability factor $\omega_g$ was assumed to equal the membrane permeability $\omega_c$. The threshold and saturation stress of MS channels were reduced from Jiang and Sun[26] to provide better agreement with our experiments (Fig 3). The leak channel permeability factor $\zeta$ was assumed to equal the threshold permeability of the MS channels (i.e. $\beta\sigma_c$). The area of the cell membrane adhered between cells $S_g$ was estimated to be on the order of 10% the reference cell surface area. As the remaining parameters could not be uniquely identified from our experiments[11], we confined them to the ranges reported by Jiang and Sun for single cell volume dynamics (as denoted by *).**

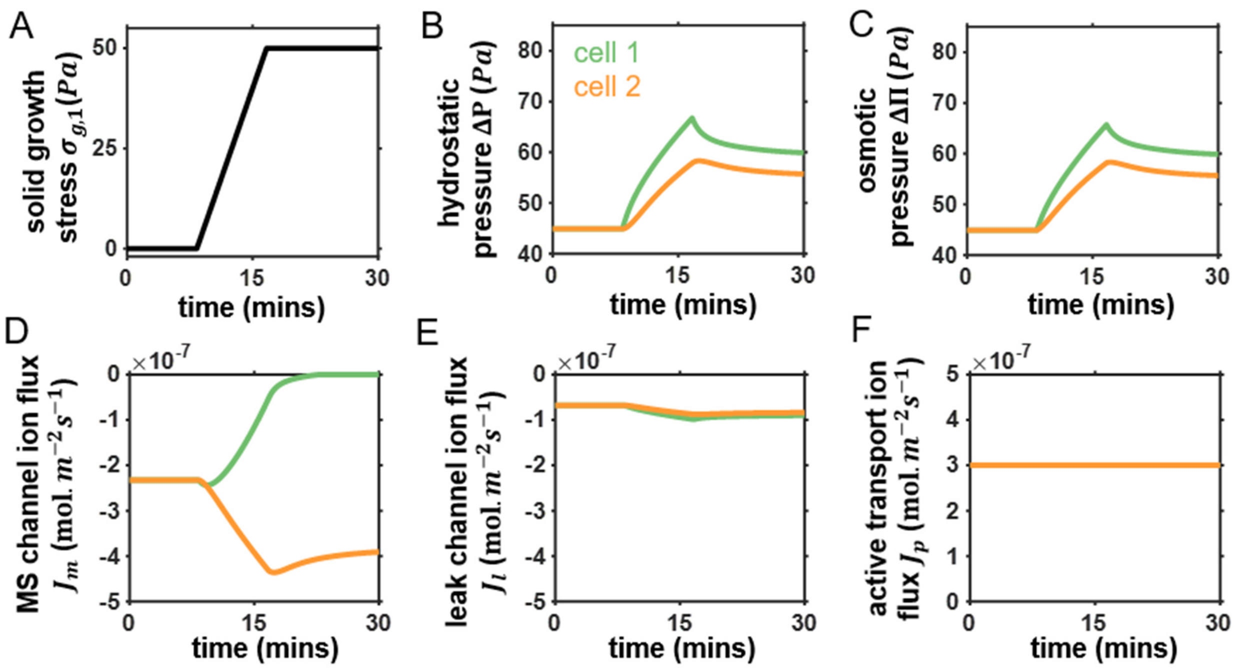


**Figure S1:** ***Additional figures for two-cell analysis under control conditions*. A) Form of solid growth stress applied to cell 1; Difference between internal and external B) hydrostatic pressure $\Delta P$ and C) osmotic pressure $\Delta\Pi$; Ion fluxes across the cell membrane: D) MS channels $J_{\mathrm{m}}$; E) leak channels $J_{\mathrm{l}}$; F) active transport $J_{\mathrm{p}}$.**

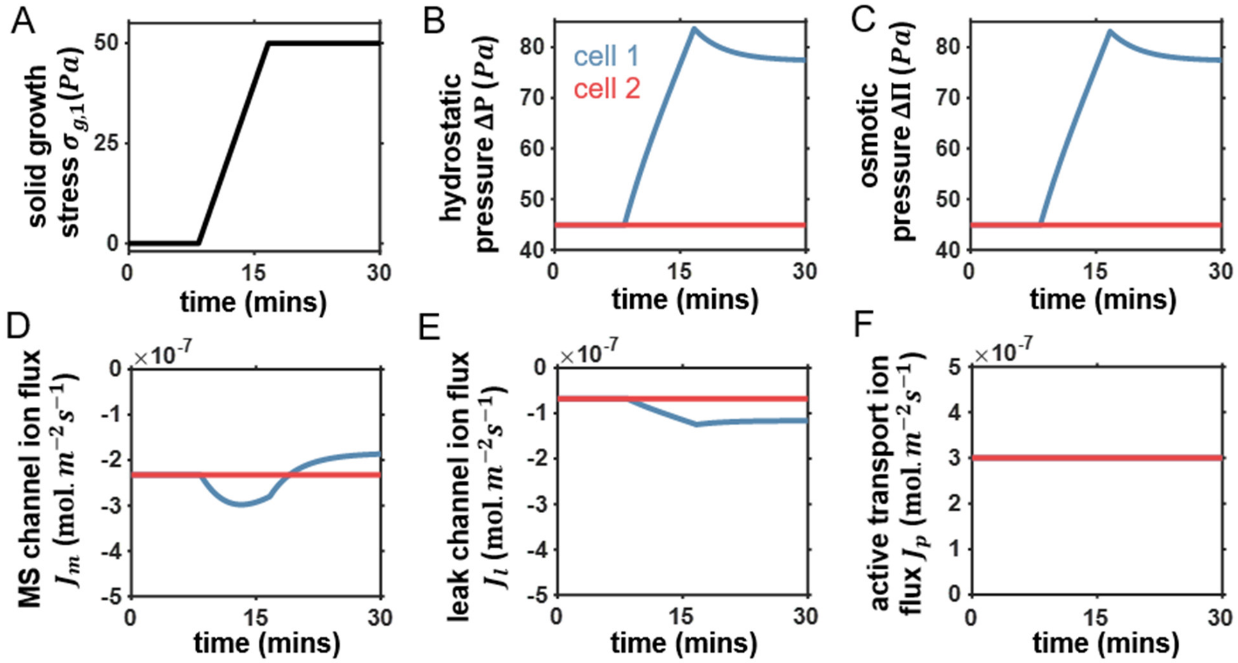


**Figure S2:** ***Additional figures for two-cell analysis during inhibition of ion flux across gap junctions*****. A) Form of solid growth stress applied to cell 1; Difference between internal and external B) hydrostatic pressure $\Delta P$ and C) osmotic pressure $\Delta\Pi$; Ion fluxes across the cell membrane: D) MS channels $J_{\mathrm{m}}$; E) leak channels $J_{\mathrm{l}}$; F) active transport $J_{\mathrm{p}}$.**

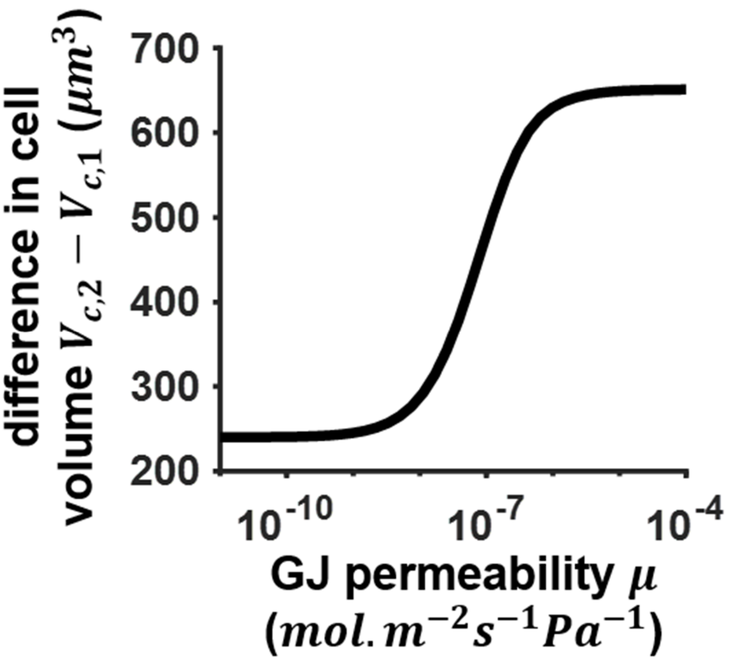


**Figure S3: Increasing gap junction permeability $\mu$ increases the steady state difference in volume $V_{c,2} - V_{c,1}$ between the 2 cells in response to a mechanical load of 50 $Pa$ on one cell.**

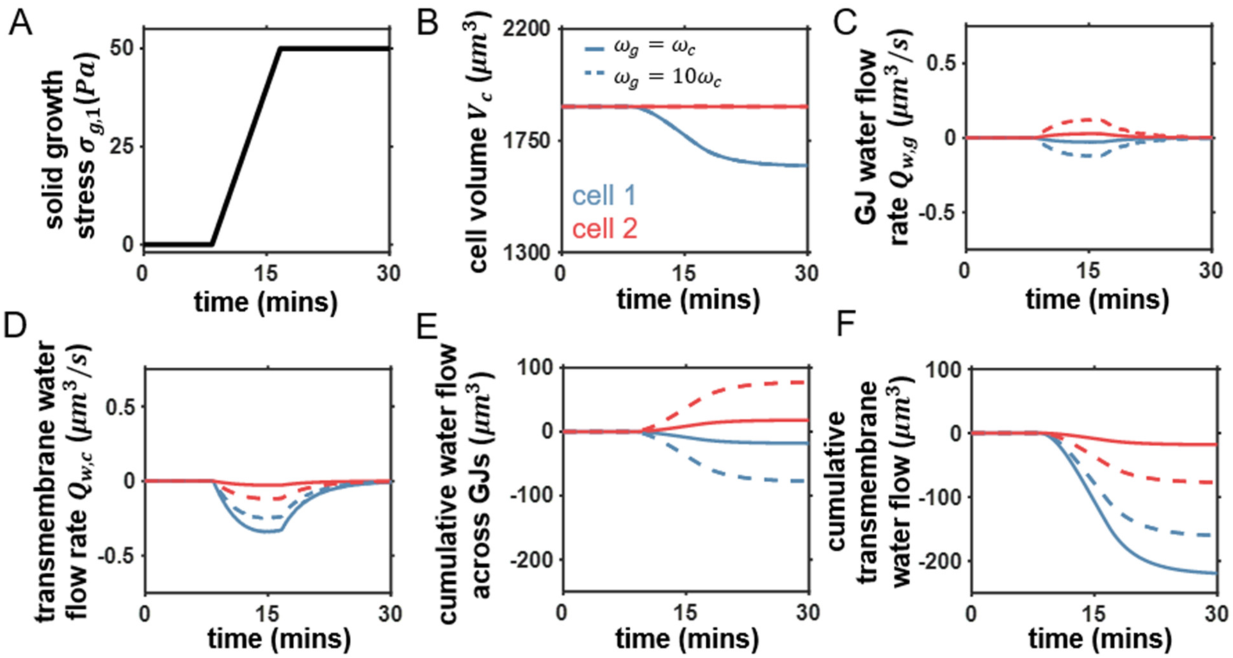


**Figure S4:** ***Ion transport inhibited across gap junctions (GJs) but water flow permitted (i.e. $\mu = 0; \omega_g > 0$).*** **A) Form of solid growth stress applied to loaded cell; For intermediate ($\omega_g = \omega_c$) and high ($\omega_g = 10\omega_c$) GJ water permeability: B) Volume change in loaded cell and its neighbor; C) GJ water flow rate $Q_w = SJ_w$; D) Transmembrane water flow rate $Q_w = SJ_w$; E) Cumulative water flow across GJs; F) Cumulative transmembrane water flow;**

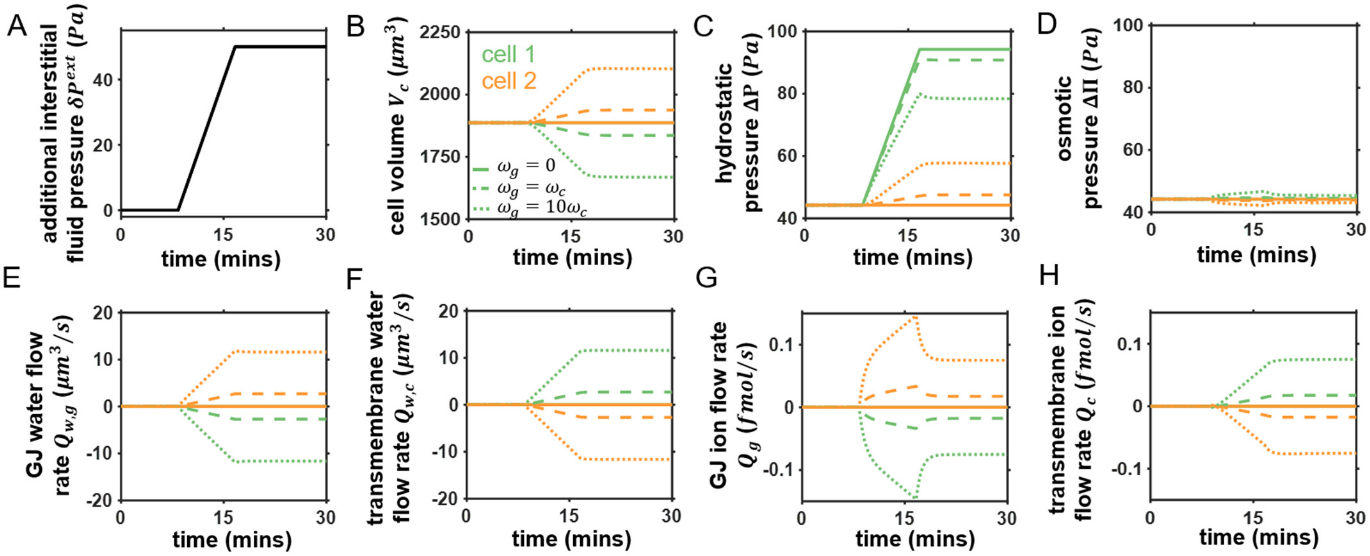


**Figure S5:** ***Cell volume change in response to increased interstitial fluid pressure*****.** **A) Form of additional interstitial fluid pressure outside cell 1; For blocked gap junctions (solid line-$\omega_g = 0; \mu = 0$), and intermediate (dashed line-$\omega_g = \omega_c$) and high (dotted line-$\omega_g = 10\omega_c$) GJ water permeability: B) Volume change in loaded cell and its neighbor; C) Hydrostatic pressure; D) Osmotic pressure; E) GJ water flow rate $Q_w = SJ_w$; F) Transmembrane water flow rate $Q_w = SJ_w$; G) GJ ion flow rate $Q = SJ$; H) Transmembrane ion flow rate $Q = SJ$; The GJ ion permeability $\mu = 5x10^{-7}\ mol.m^{-2}s^{-1}Pa^{-1}$ unless otherwise stated.**

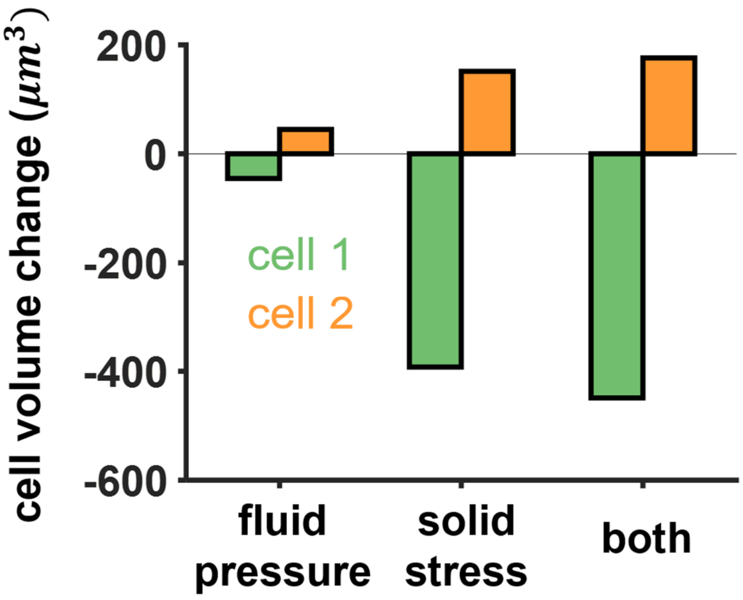


**Figure S6: Cell volume change at steady state in response to increased interstitial fluid pressure or solid stress on cell 1. Fluid pressure $\delta P_1^{ext} = 50Pa$; solid stress $\sigma_{g,1} = 50Pa$; both $\delta P_1^{ext} = \sigma_{g,1} = 50Pa$.**

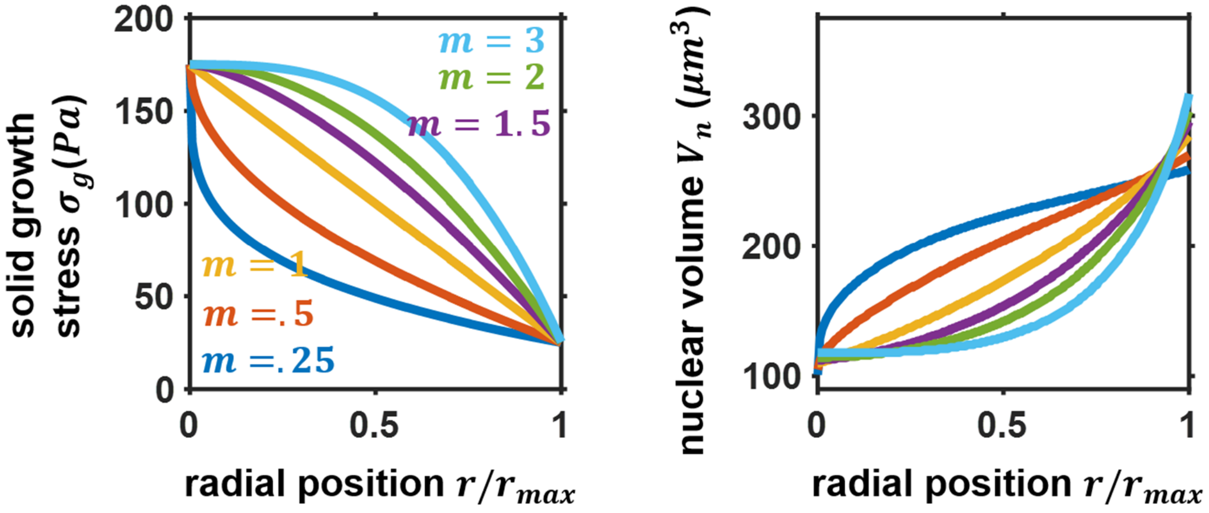


**Figure S7: Influence of alternative loading profiles on spatial cell volume in the simulated cancer organoid. Solid growth stress $\sigma_g(r) = \delta\sigma_g^{max} - \left(\delta\sigma_g^{max} - \delta\sigma_g^{min}\right)\left(\frac{r}{r_{max}}\right)^m$. In all simulations $\delta\sigma_g^{min} = 25\ Pa$ unless otherwise stated.**

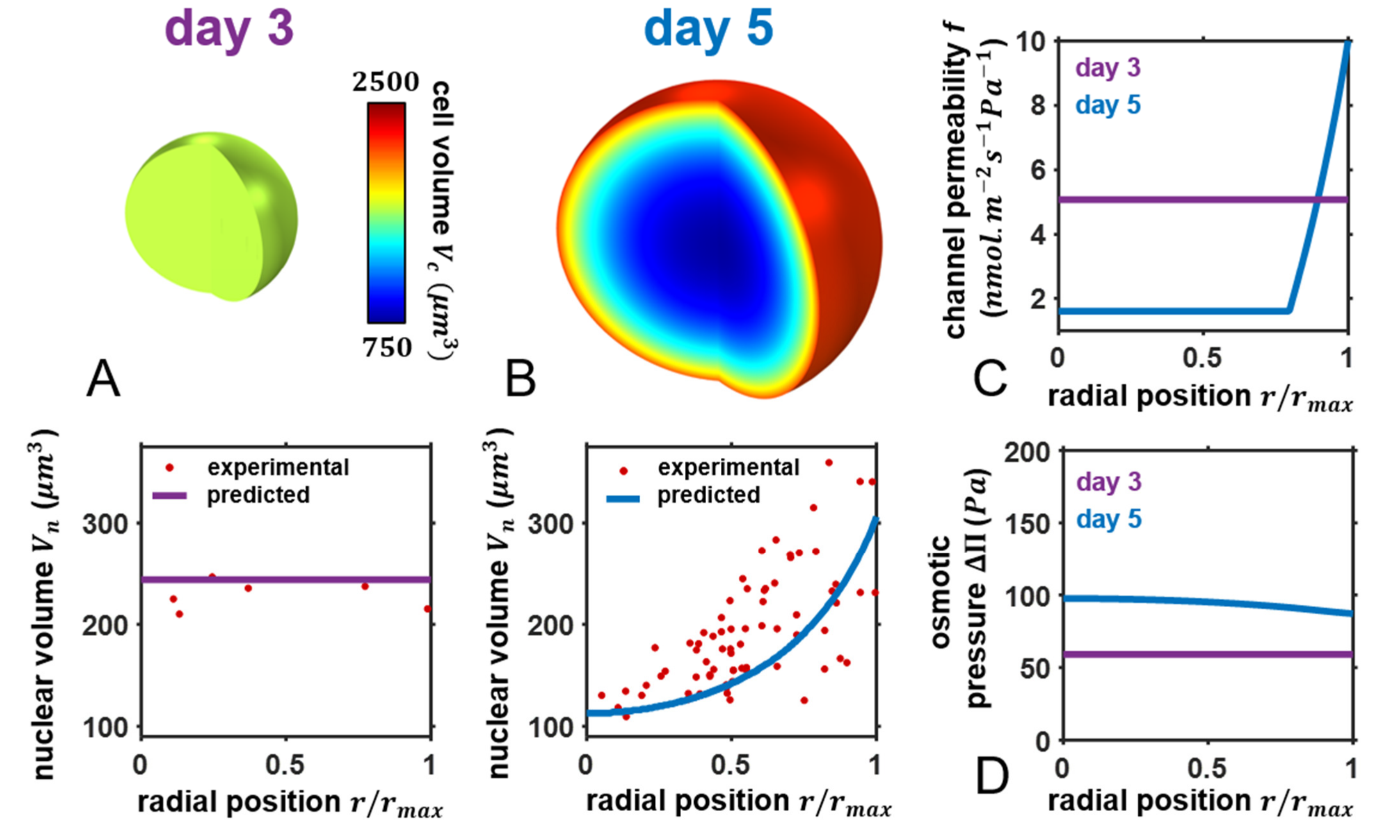


**Figure S8: A) Predicted and experimental spatial cell and nuclear volumes under control conditions on A) day 3 and B) day 5; Predicted C) spatial channel permeability $f' = f(\sigma) + \zeta$ and D) osmotic pressure $\Delta\Pi$. Solid growth stress on day 3 $\sigma_g(r) = 25\ Pa$.**

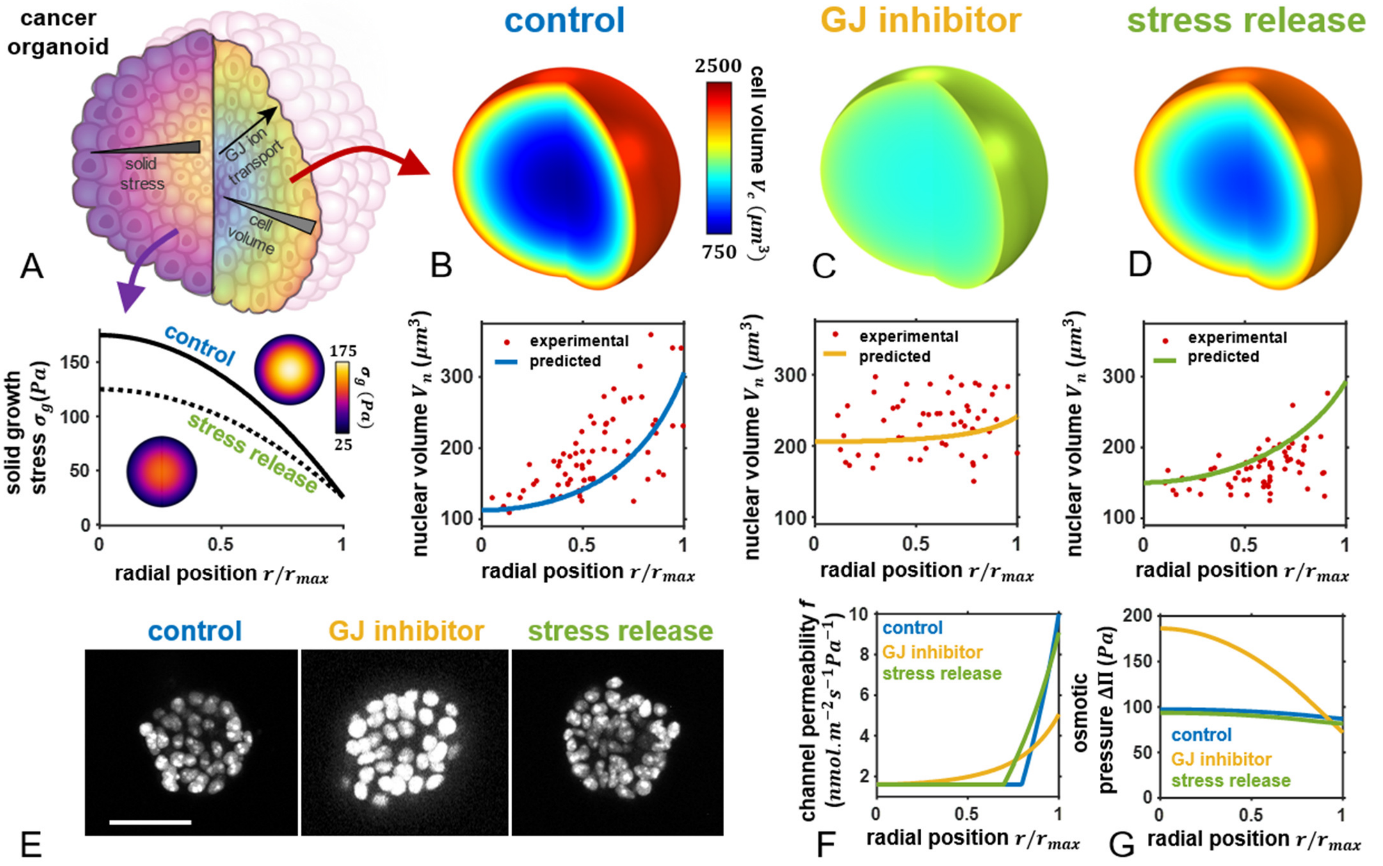


**Figure S9:** ***Spatially non-uniform cell volume in cancer organoid*.** **A) Solid growth stress $\sigma_g$ is spatially non-uniform; Predicted and experimental (day 5) spatial cell and nuclear volumes under B) control, C) inhibited gap junction, and D) stress release conditions; E) Cross-section images of GFP-NLS-labelled MCF 10A cells at day 5. Scale bar 50 $\mu m$; Predicted F) spatial channel permeability $f' = f(\sigma) + \zeta$ and G) osmotic pressure $\Delta$**

**SI Note 1: Analytical solutions at limits of gap junction permeability**

From our continuum formulation, it is possible to analytically derive solutions at the limits of gap junction permeability. Considering again the expression for the number of cellular ions at position $r$:

$$\frac{\partial n(r)}{\partial t} = S_g \mu \frac{r_0^2}{r^2}\frac{\partial}{\partial r}\left(r^2 \frac{\partial \Delta\Pi(r)}{\partial r}\right)$$
$$-4\pi r_c^2(r)\Big(\big(f(\sigma(r)) + \zeta + \gamma\big)\Delta\Pi(r) - \gamma\Delta\Pi_c\Big), \qquad (S1)$$

At steady state, $\frac{\partial n(r)}{\partial t} = 0$, thus considering the limit of zero gap junction permeability (i.e. $\mu = 0$) yields:

$$\Delta\Pi(\mathrm{r}) = \frac{\gamma\Delta\Pi_c}{f(\sigma(r)) + \zeta + \gamma}. \qquad (S2)$$

As at steady state $\Delta\Pi(\mathrm{r}) = \Delta P(r)$ (via Eqn 3), mechanical equilibrium dictates:

$$\sigma(\mathrm{r}) = \frac{K}{2}\left(\frac{r_c^2(r)}{r_0^2} - 1\right) + \sigma_a$$
$$= \frac{\big(\Delta\Pi(r) - \sigma_g(r)\big) r_c(r)}{2h}. \qquad (S3)$$

Therefore, spatial cell volume can obtained for a given solid stress distribution $\sigma_g(r)$.

Next, considering the limit of infinite gap junction permeability ($\mu = \infty$) mandates that $\frac{\partial}{\partial r}\left(r^2 \frac{\partial \Delta\Pi(r)}{\partial r}\right) = 0$. As such, $\Delta\Pi(\mathrm{r})$ must be constant and spatially uniform (i.e. $\Delta\Pi(\mathrm{r}) = \Delta\Pi^*$). We can then multiply Eqn S1 by $r^2$ and integrate between 0 and $r_{max}$ to obtain:

$$4\pi \int_0^{r_{max}} r^2\, r_c^2(r)\big(f(\sigma(r)) + \zeta + \gamma\big)\Delta\Pi^* dr$$
$$= 4\pi(\Delta\Pi_c\gamma)\int_0^{r_{max}} r^2\, r_c^2(r) dr, \qquad (S4)$$

which leads to:

$$\Delta\Pi^* = \frac{\gamma\Delta\Pi_c}{\gamma + \frac{< f(\sigma(r))\, r_c^2(r) >}{< r_c^2(r) >}}, \qquad (S5)$$

where $< r_c^2(r) > = \int_0^{r_{max}} r^2\, r_c^2(r) dr$

and $< f(\sigma(r))\, r_c^2(r) > = \int_0^{r_{max}} f(\sigma(r))\, r^2\, r_c^2(r) dr$.
